# Smoothened turnover regulated by Hedgehog signaling in Drosophila

**DOI:** 10.64898/2026.01.08.698469

**Authors:** Ryo Hatori, Wanpeng Wang, Thomas B. Kornberg

## Abstract

During development, the processes of cell-cell signaling orchestrate growth and morphogenesis using both static features and moving parts. For example, expression domains of genes which set states of determination may be fixed in space for periods of time, but multiple signals move between the determined cells and intracellular movements of components of the signal transduction machines are key to information exchange. In the Drosophila wing imaginal disc, Hedgehog (Hh) protein produced in the posterior compartment moves to cells across the anterior/posterior compartment border to induce signal transduction and various intracellular responses. In this work, we investigated protein turnover and intracellular trafficking of components of Hh signal transduction in the wing disc and found that both are major features of Hh signaling.

**Summary:** The intracellular distribution and half-life of Smoothened is regulated by Hedgehog signaling.

## Introduction

Hedgehog signaling regulates cell fate choices, morphogenesis, and patterning during animal development, and in the human, several types of cancers are associated with its mis-regulation (Ingham, 2018). Hh signaling in the Drosophila wing imaginal disc involves Hh production by posterior (P) compartment cells and Hh signal transduction in anterior compartment (A) cells across the A/P compartment border. Levels of Hh in A compartment cells decline with increasing distance from the border and correlate with a declining gradient of Hh signal transduction. Hh uptake and signaling in A cells engage three transmembrane proteins which are core pathway components: co-receptors Patched (Ptc) and Interference hedgehog (Ihog), and the seven-pass transmembrane protein Smoothened (Smo).

The epithelial cells of the wing disc are columnar, with an average diameter of 2.5 μm and length that extends 40-50 μm along their apicobasal axis. Although Hh secretion by P cells and uptake by A cells is basolateral (Callejo et al., 2011; Chen et al., 2017), the relatively small apical compartment in these cells - defined basally by the adherens and septate junctions <5 μm from the apical surface - has an important role in Hh signaling. Hh translocates to the apical plasma membrane after it is synthesized, but it is not released apically. Instead, it is internalized by dynamin-dependent endocytosis to endosomal vesicles that move to basolateral membranes and basolateral cytonemes that extend from Hh-producing cells toward the A compartment. (Callejo et al., 2011). Hh also moves along basolateral cytonemes that extend from A compartment Hh-receiving cells where it is visible colocalized with Ptc (Chen et al., 2017). Genetic conditions that either perturb the basolateral cytonemes or the apical to basolateral choreography of Hh in producing cells compromise Hh dispersion and the Hh concentration gradient in the A compartment (Callejo et al., 2011; Chen et al., 2017; D’Angelo et al., 2015).

Activation of the Hh pathway in A compartment wing disc cells changes levels and subcellular locations of several signal transduction components. *ptc* RNA is expressed specifically in the A compartment and at low levels except in the cells near the A/P compartment border in which expression is upregulated by Hh signaling. Hh signaling also induces the redistribution of Ptc protein from the plasma membrane to intracellular vesicles, and changes the form, expression levels and subcellular location of the Cubitus interruptus (Ci) transcription factor (Aza-Blanc et al., 1997). Similar to *ptc*, *ci* expression is specific to A compartment cells and Ci levels are upregulated by Hh signaling in cells near the compartment border. In the absence of Hh signaling, Ci is proteolyzed and converted to a transcriptional repressor that translocates to the nucleus from the cytoplasm. Hh signaling blocks Ci proteolysis and blocks Ci conversion to its repressor form. A cells that activate Hh signaling at the compartment border are characterized by elevated levels of cytoplasmic Ptc and Ci.

Smo is an activator of the pathway that is regulated by Hh in several ways (Denef et al., 2000). In the absence of Hh, Ptc inhibits Smo, but Hh signaling relieves this inhibition. Smo is also regulated post-translationally and by intracellular trafficking. Although *smo* RNA is expressed uniformly in the A and P compartments, Levels of Smo protein are high in P cells, decline in A cells near the A/P compartment border and drop to lower amounts further from the border. The reduced levels of Smo in A compartment cells are regulated by Hh signaling. Hh signaling also induces Smo to relocate from cytoplasmic stores to the basolateral plasma membrane (Zhu et al., 2003). Blocking endocytosis, for example by expression of dominant negative Rab7 or dominant negative dynamin, results in basolateral accumulation of Smo and upregulation of downstream target genes (Zhu et al., 2003). In contrast, blocking endosomal sorting results in accumulation of Smo in intracellular vesicles and pathway deactivation (Jiang et al., 2018).

These effects highlight the importance of understanding how Smo levels and intracellular distributions are regulated. However, despite extensive characterizations of Smo (Kowatsch et al., 2019), the spatiotemporal dynamics that link Hh uptake to Smo distributions and to Hh target gene activation remain unknown. In this study, we used fluorescent tags to monitor levels and intracellular distributions of Hh and Smo.

## Results

### Contrasting intracellular distributions of Hh in A and P compartment cells

To investigate possible causal relationships between Hh, Ptc and Smo distributions in cells that transduce Hh signaling, we first compared the relative levels of Hh in A and P compartment cells to obtain a measure of Hh involved in signal transduction. Whereas all P cells appear to express Hh RNA and protein, A cells do not have detectable levels of *hh* RNA, and only the A cells near the A/P compartment border have detectable levels of Hh protein (Tabata and Kornberg, 1994). To estimate the relative amounts of A and P Hh, we compared the levels of Hh:GFP in discs obtained from larvae whose only *hh* gene is a BAC transgene. This transgene includes approximately 40kb of genomic sequence surrounding a *hh* transcription unit into which GFP coding sequences are inserted. Animals with this single *hh* transgene grow normally and are indistinguishable from wild type. The measured ratio of GFP was approximately 95% P and 5% A (Fig. 1A-D). If Hh turnover is similar in A and P cells, this result is consistent with the idea that most Hh does not exit producing cells and that the amount of Hh that activates signal transduction is small relative to total production.

**Figure 1.**
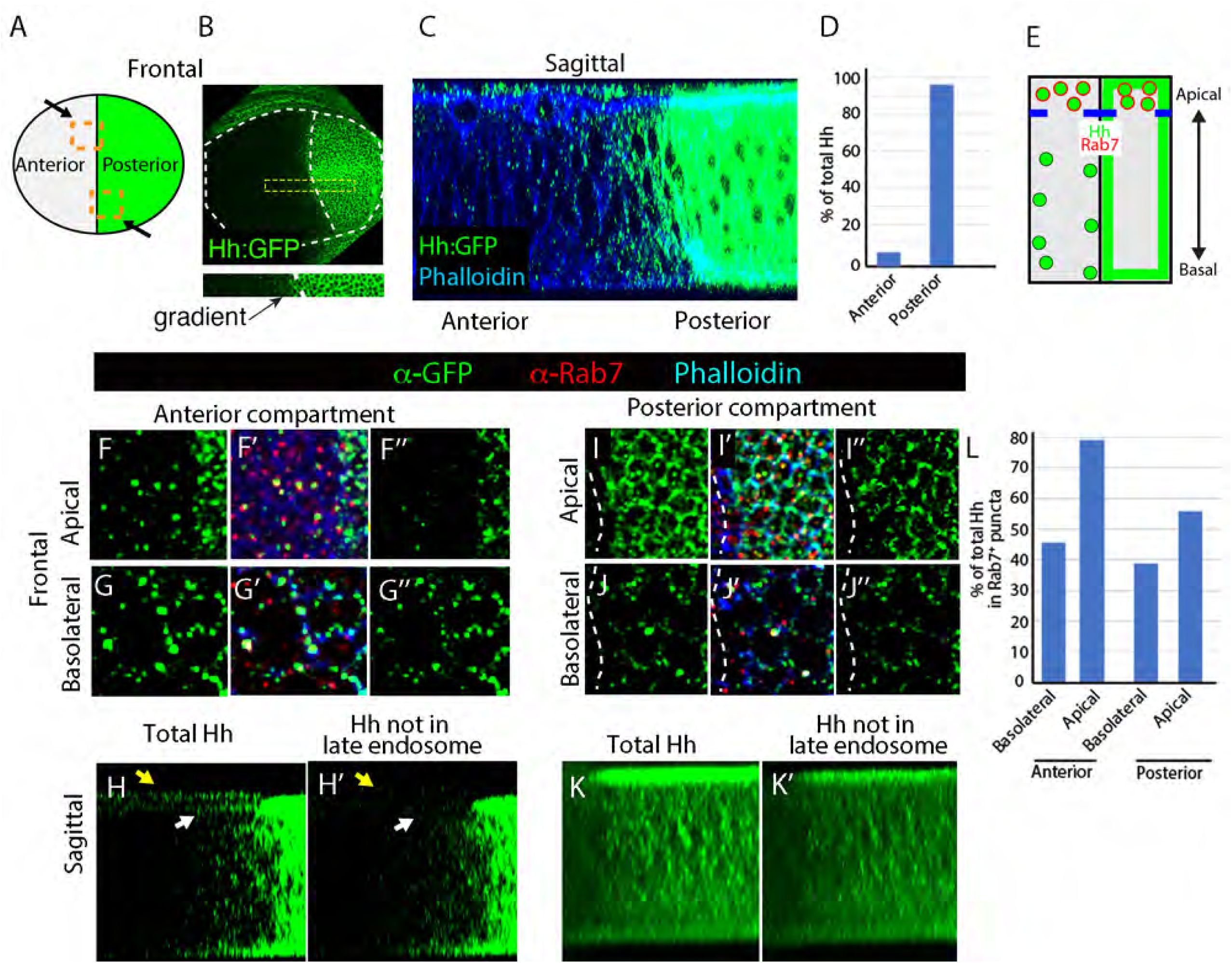
Subcellular Hh distributions in the wing disc. (A) Schematic of frontal depiction of the wing pouch of the wing disc with the A (gray) and P (green) compartments indicated by gray and green, respectively; boxes indicate positions of (F-K) images. (B) White dotted lines delineate wing pouch and compartment border of disc expressing BAC encoded Hh:GFP. (C) Sagittal section of the wing disc expressing BAC encoded Hh:GFP (α-GFP antibody staining). Septate junction stained with α-DLG (red) marks septate junction; phalloidin marks adherens junction (blue). (D) Graph of Hh:GFP fluorescence in A and P compartments. (E) Schematic of sagittal wing pouch section with apical and basolateral compartments, Hh:GFP (green) and Rab7 (red) indicated. (F-G”) Frontal views of region of the A compartment of wing discs with BAC-encoded Hh:GFP detected with α-GFP antibody (green) and stained with α-Rab7 antibody (red) and Phalloidin (blue). (H,H’) Sagittal section showing total Hh signal (H) and Hh not colocalized with Rab7. White and yellow arrows indicate peripodial membrane and apical compartment, respectively. (I-K’) Same as (F-H’) for P compartment. (K) Graph showing the proportion of BAC encoded Hh:GFP signal that overlaps with Rab7 in apical/basolateral optical sections.

**Figure 1S.**
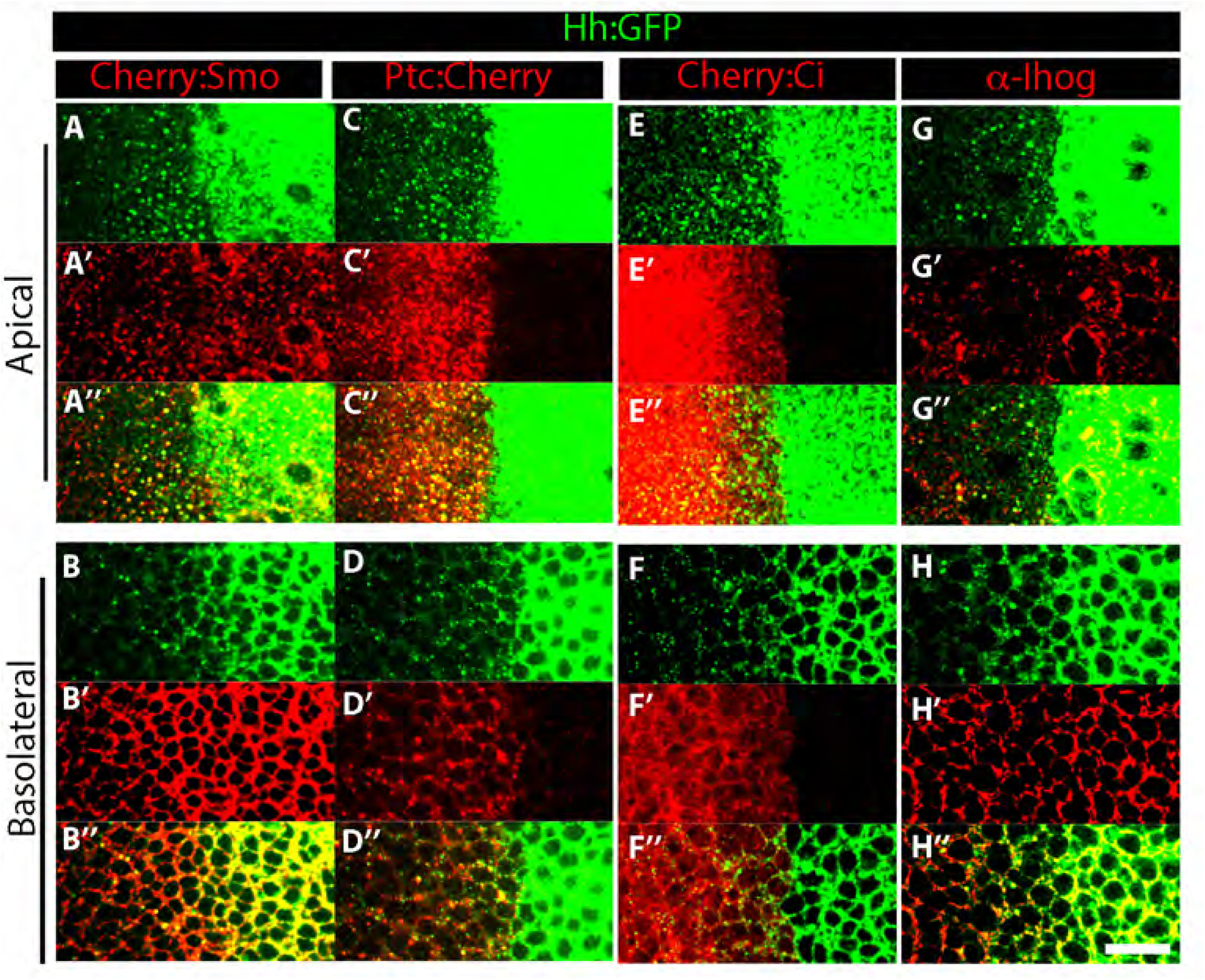
Apicobasal distributions of Hh signaling pathway components. Apical and basolateral optical sections of wing discs. BAC-encoded Hh:GFP (green), Cherry (red). (A-B) BAC-encoded Cherry:Smo; (C-D) BAC-encoded Ptc:Cherry; (E-F) BAC-encoded Cherry:Ci ;(G-H) α-Ihog (red).

To investigate Hh distributions in more detail, we examined the subcellular localizations of BAC-encoded Hh:GFP as well as the Rab7 GTPase, a marker of late endosomes. Hh was present in intracellular puncta in both A and P cells, and at the plasma membrane in P cells (Fig. 1E). Hh:GFP in A cells was detected only in intracellular puncta in both the apical and basolateral compartments (Fig. 1E-H). The majority of apical Hh in the A compartment colocalized with Rab7, suggesting that most apical Hh is in the process of degradation in late endosomes (Fig. 1L). In Hh-producing P cells, most basolateral Hh was associated with the plasma membrane, while apical Hh was both membrane-associated and in intracellular puncta. Approximately half apical Hh colocalized with Rab7, suggesting that this Hh has entered the endosomal pathway and is destined for degradation (Fig. 1I-L). The apical Hh that is vesicular but not Rab7-associated may include Hh that is destined to be transported to basolateral compartments for dispersion, as proposed by Callejo et al., 2011 (Callejo et al., 2011). These distributions are consistent with the idea that Hh enters A cells through their basolateral compartment (Callejo et al., 2011) and then moves apically after activating signal transduction (Kornberg, 2014).

To investigate whether Hh might affect the localization of A cell signal transduction components, we also examined Ptc, Smo, Ihog, and Ci distributions. We observed higher levels of Cherry:Smo and Ptc in cells adjacent to the compartment boundary that receive Hh compared to cells further away from the compartment boundary (Fig. S1A-D). These Smo and Ptc distributions are Hh dependent (Denef et al., 2000). Based on relative fluorescence, the amount of Hh in these cells was significantly lower than Ptc or Smo, suggesting that the effects of Hh on Smo and Ptc are not stoichiometric. Most Smo, Ptc, and Ihog fluorescence was punctal apically and membrane associated in the basolateral regions (Fig. 1SA-D, G-H). The distribution of Ci was uniform in both apical and basolateral regions (Fig. 1SE-F). Relative levels of Ihog were higher in the basolateral membrane compared to apical membrane (Fig. 1SG-H).

### Live fluorescence of GFP and Cherry fusion proteins identifies distinct pools of Smo

To investigate the effects of Hh signaling in the basolateral region, we focused on Smo, because its basolateral localization correlates with signal transduction activity (Gonçalves Antunes et al., 2022). Previously, we generated transgenic Drosophila lines that express physiological levels of GFP tagged Smo from a BAC transgene (Chen et al., 2017). For this study, we also generated BAC encoded Cherry:Smo and a form with tandem tags, GFP:Cherry:Smo (Fig. 2A). These BAC encoded fusion proteins rescued the lethality of the *smo^3^* allele (Chen and Struhl, 1998).Wings of the rescued flies had morphologically normal wings (Fig. 2B-D), indicating that the fusion Smo constructs are functional.

**Figure 2.**
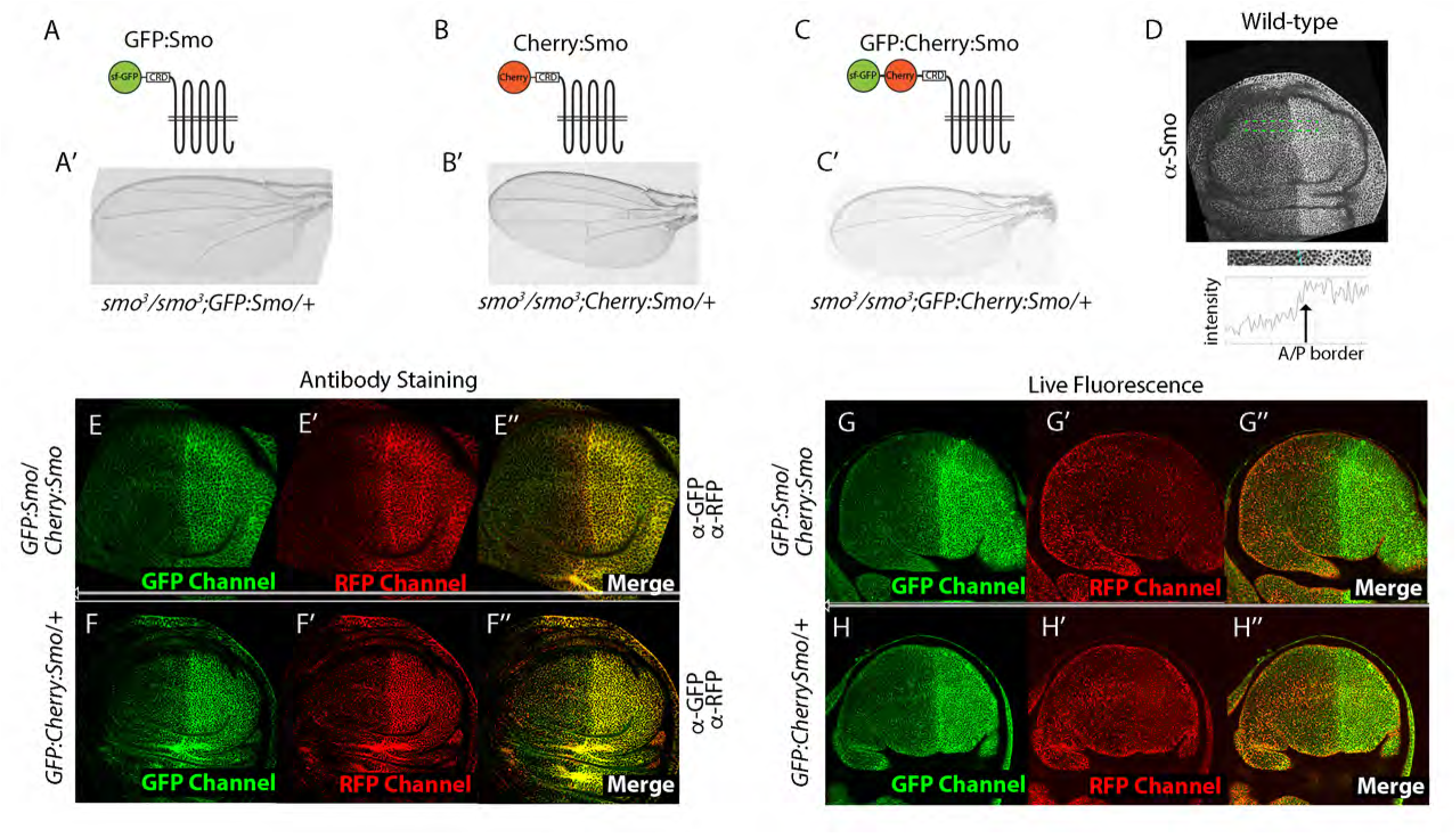
Contrasting patterns of GFP:Smo and Cherry:Smo fluorescence in wing discs. (A-C) Schematic diagrams of GFP:Smo, Cherry:Smo, and GFP:Cherry:Smo fusion proteins and *smo^3^*wings from flies grown at 25°C with BAC-GFP:Smo (A’), BAC-Cherry:Smo (B’), and BAC-GFP:Cherry:Smo (C’) transgenes. (D) Wing disc from wild type fly stained with α-Smo antibody. Intensity plot of area bounded by green dashed lines below, with location of compartment border indicated. (E-H”) Frontal optical section images of discs with genotypes *GFP:Smo/Cherry:Smo* (E,G) and *GFP:CherrySmo/+* (F,H) acquired with GFP (Green) or RFP (Red) channel. (E-F”) Fixed and stained with a-GFP or a-RFP antibodies. (G-H”) Unfixed, live fluorescence.

Using these Smo fusion proteins, we first compared the distributions of BAC-encoded GFP:Smo, Cherry:Smo and GFP:Cherry:Smo by applying antibodies directed against GFP and RFP to fixed wing discs (Fig. 2E-F). In frontal sections imaged at basolateral optical planes,:GFP and Cherry fluorescence levels were higher in P compared to the A cells, and the relative levels of GFP and Cherry fluorescence were similar in both compartments (Fig. 2E,F). Most of the fluorescence appeared to be membrane-associated. These distributions are similar to the distributions of Smo detected by anti-Smo antibody (Fig. 2D).

We also imaged GFP:Smo and Cherry:Smo fluorescence in unfixed, live wing discs and found that the levels and subcellular distributions of GFP and Cherry fluorescence differed (Fig. 2G,H). Whereas most of the GFP fluorescence appeared to be membrane-associated, Cherry fluorescence was punctate and not apparently membrane associated. GFP fluorescence levels were higher in P cells compared to A cells, a pattern similar to the Smo distribution detected by anti-Smo antibody. In contrast, Cherry fluorescence was punctate, more evenly distributed between the A and P compartments, and at relatively low levels. These results suggest that the GFP and Cherry fluorescence of GFP:Smo, Cherry:Smo, and GFP:Cherry:Smo identify different pools of Smo in live cells.

### Cherry fluorescence identifies Smo in a degradation pathway

GFP and Cherry have different pH sensitivities: GFP is more sensitive to acid pH such that the absence of GFP fluorescence might be evidence of an acidic quenching. To test if the differences in GFP and Cherry fluorescence in unfixed wing discs might be a consequence of localization to acidic compartments, we treated discs with ammonium chloride, a condition that has been used to neutralize acidic compartments in neurons (Boron and De Weer, 1976). We compared GFP and Cherry fluorescence of GFP:Cherry:Smo in control discs and in discs treated with ammonium chloride. In the A compartment of control discs, cells had many puncta with only Cherry fluorescence. In contrast, most fluorescent puncta had both GFP and Cherry fluorescence in discs incubated with ammonium chloride (Fig. 3A,B). The Cherry fluorescence of GFP:Cherry:Smo may therefore identify Smo in acidic compartments. GFP fluorescence in the P compartment was not apparently changed by the ammonium chloride treatment.

**Figure 3.**
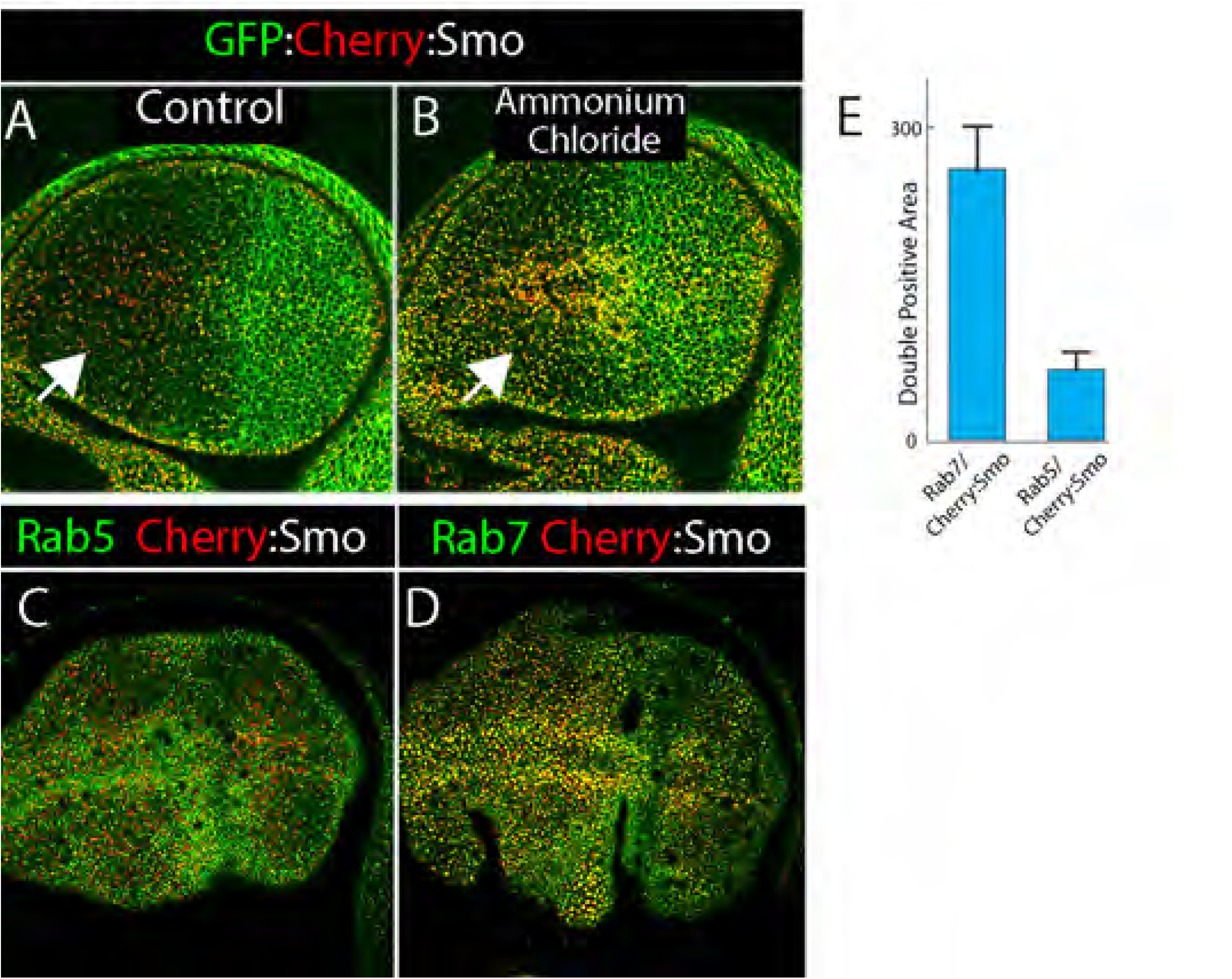
Cherry:Smo fluorescence localizes to late endosomes. (A-E) Frontal views of unfixed wing discs, optical section close to the apical surface; left, anterior; right, posterior. (A,B) Wing discs expressing BAC-encoded GFP:Cherry:Smo without (A) and with ammonium chloride (B); GFP channel (green), RFP channel (red). (C) Wing disc expressing BAC-encoded GFP:Smo (green) and Rbcn3A RNAi (expressed during L3 at 29°C) in the dorsal region (below white dashed line) using (*ap-Gal4/tub-Gal80^ts^; GFP:Smo/Rbcn3A-RNAi*). (D,E) Unfixed wing discs expressing Cherry:Smo (red) and Rab5:YFP (green) (D) or Cherry:Smo (red) and Rab7:YFP (green). (F) Amount of Cherry:Smo/Rab7 double positive and Cherry:Smo /Rab5 double positive puncta.

Because acidic compartments are associated with degradation and endocytic compartments, GFP:Cherry:Smo with only Cherry fluorescence may represent a Smo pool in the degradation pathway. To test this idea, we imaged Cherry:Smo together with Rab5 and Rab7. Rab5 is a marker for early endosomes, and Rab7 is a marker for late endosome (reviewed in (Stenmark, 2009)). Whereas Cherry:Smo was not observed to be co-localizeed with Rab5, co-localization with Rab7 was frequent (Fig. 3D-E). These results are consistent with the idea that Cherry:Smo fluorescence marks Smo that in the process of degradation.

### Apical-basal localization of Smo

We next investigated the subcellular locations of the two Smo pools. To characterize the subcellular distributions of Smo in wing disc cells, we first characterized total Smo protein by staining wing discs with anti-Smo antibody in animals that express Smo endpgenously. Staining fixed discs with anti-Smo antibody revealed both apical and basolateral Smo; most protein was basolateral. In contrast, unfixed discs with the GFP:Smo and Cherry:Smo BACs had Cherry fluorescence that was mostly apical and GFP fluorescence that was almost exclusively basolateral (Fig. 4B,B’). To characterize the apical-basal localization of GFP and Cherry fluorescence in more detail, we imaged the GFP:Smo and Cherry:Smo proteins together with markers of the adherens junction. Adherens junctions form a ring around epithelial cells at their apical/basolateral boundary and connect cells to their neighbors (Fig. 1B). Imaging the adherens junction component E-Cadherin (E-Cad) together with GFP:Smo revealed that GFP fluorescence was basal to E-Cad in both A and P compartments (Fig. 4C-G). In contrast, Cherry fluorescence was mostly at the level of or more apical to E-Cad in both compartments (Fig. 4H-L).

**Figure 4.**
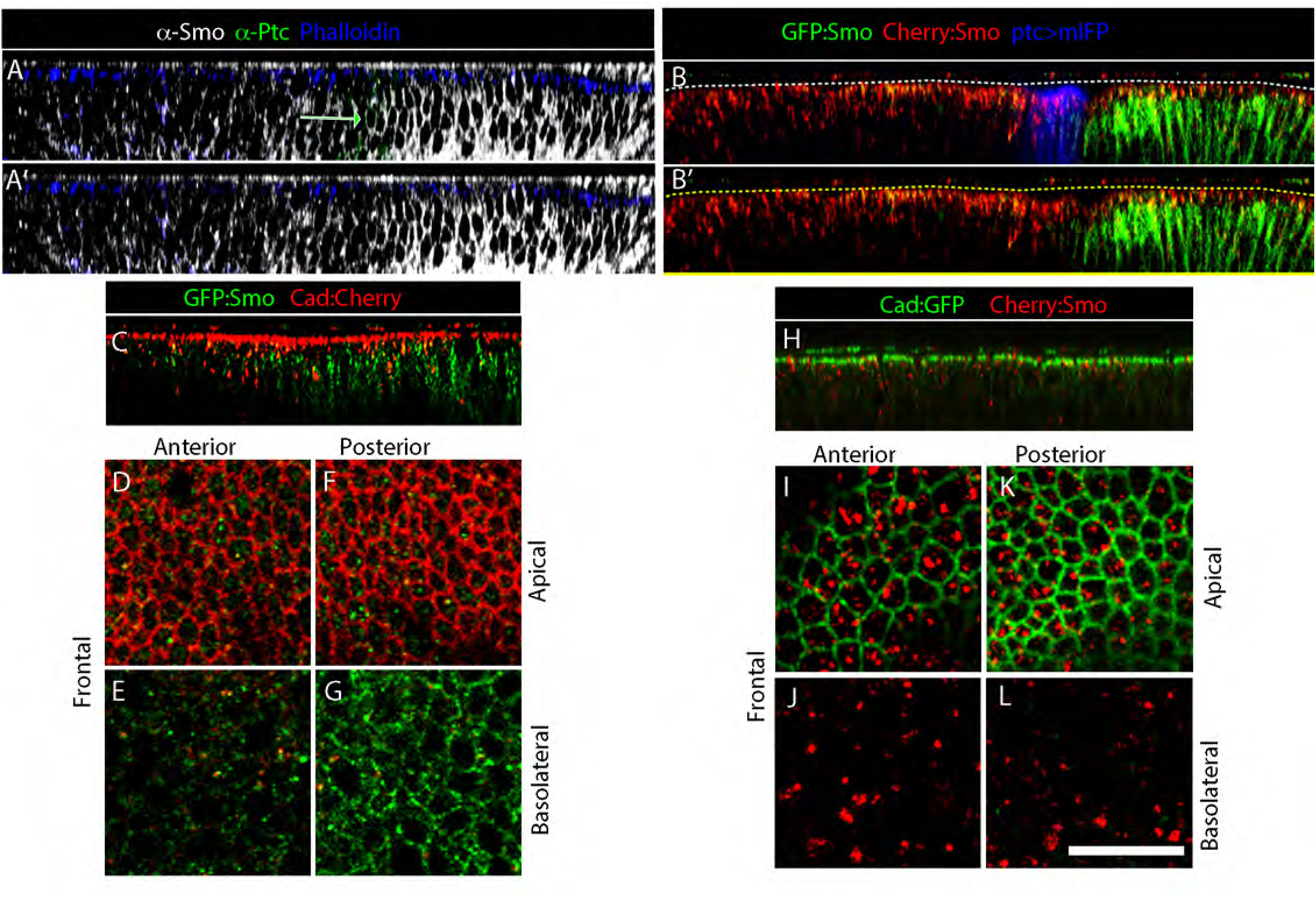
Apicobasal distribution of GFP:Smo and Cherry:Smo fluorescence. (A-C) Sagittal views of wing discs stained with (A,A’) α-Smo (gray), α-Ptc (green), and phalloidin (blue), or (B,B’) expressing BAC-GFP:Smo (green), BAC-Cherry:Smo (red), and mIFP (*ptc-Gal4>mIFP; Smo:GFP:Cherry/+*); dashed lines indicate apical surface. (C-G) Wing disc expressing BAC-GFP:Smo (green) and Cad:Cherry; (C) sagittal, (D-G) frontal views in A compartment, P compartment and in apical or basolateral sections. (H-L) Same as (C-G), except wing discs express Cad:GFP (green) and BAC-encoded Cherry:Smo (red).

Because GFP matures to become fluorescent more rapidly than Cherry, we investigated whether the BAC-encoded GFP:Smo and Cherry:Smo fusion proteins could be analyzed as fluorescent protein timers (Khmelinskii et al., 2012) to distinguish older protein from more recently produced protein. This idea would be consistent with Smo an intracellular choreography process in which Smo protein first situates basally before moving apically.

### Hh regulates the kinetics of Smo movement independently of Ptc

To investigate effects of Hh on Smo levels, we analyzed GFP and Cherry fluorescence of GFP:Cherry:Smo in wing disc cells that have different levels of Hh. Cells in the P compartment have high levels of Hh, A cells near the A/P border have lower levels, and levels in A cells farther from the border are not detectable (Fig. 1A,C). Anti-Smo antibody located Smo in the basolateral regions of both P compartment cells and in A compartment cells adjacent to the compartment boundary (Fig. 5A,B). The pattern of GFP fluorescence in GFP:Smo-expressing unfixed discs (Fig. 5C,D) was comparable to the distribution detected by antibody, with high levels throughout the P compartment and lower levels in the band of 3-4 cells adjacent to the compartment boundary. These distributions are consistent with the idea that the basolateral localization of newly synthesized Smo correlates with high levels of Hh. GFP fluorescence was not detected close to the apical surface anywhere in the disc.

**Figure 5.**
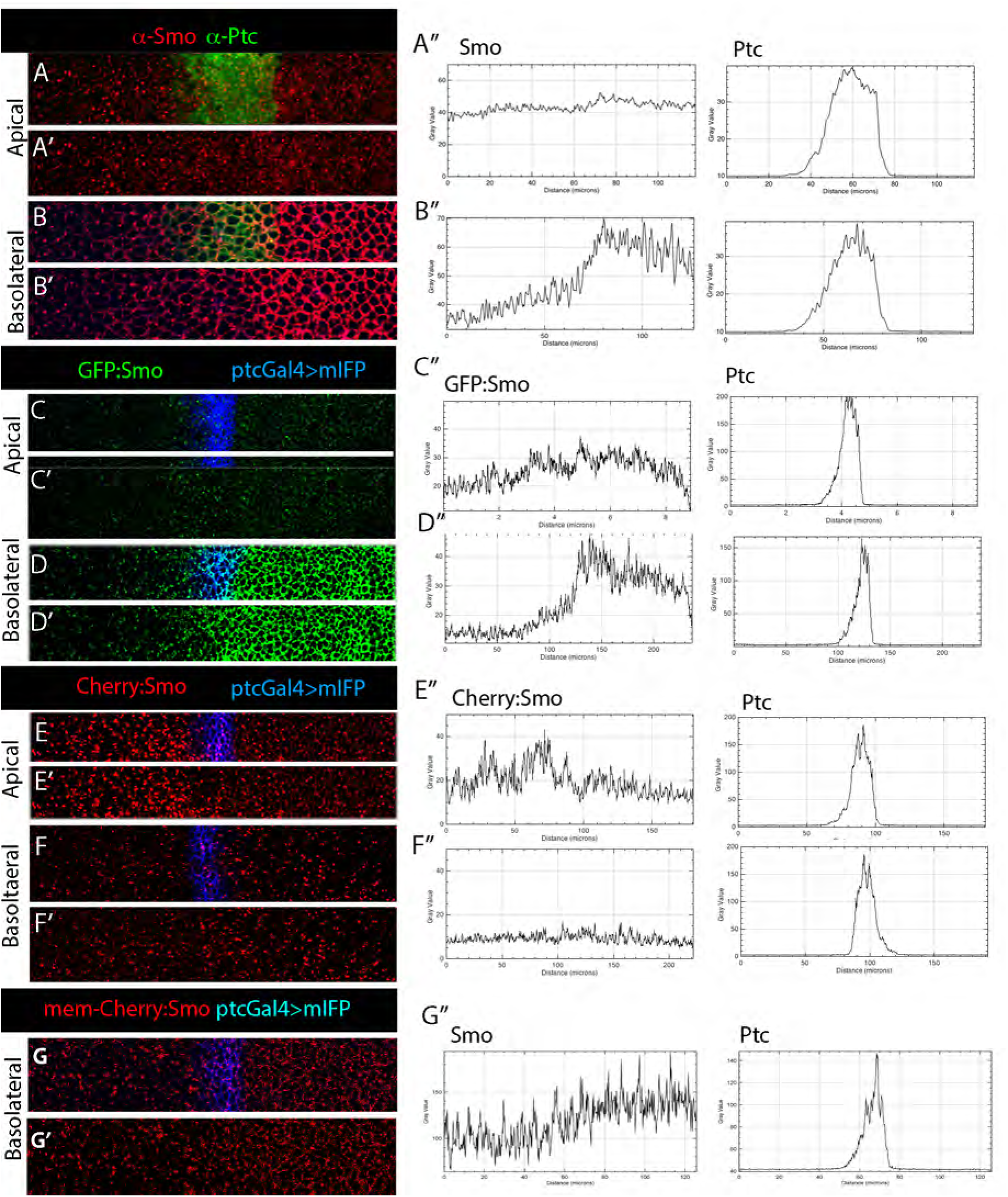
Smo kinetics and turnover regulated by Hh. Wild-type wing disc stained with α-Smo (red) and α-Ptc (green) at apical (A,A’) and basolateral (B,B’) optical planes. (A,B) Smo and Ptc staining; (A’,B’) Smo staining only. (A”,B”) Graphs showing intensity plots of images in (A,B) across A/P axis. (C,D) Same as (A,B) for unfixed disc expressing BAC-encoded GFP:Smo and mIFP driven by *ptc-Gal4* showing fluorescence of Smo (green) and mIFP (blue) (*ptc-Gal4>mIFP*; GFP:*Smo* /*+*). (E,F) Same as (C,D) for disc expressing BAC-encoded Cherry:Smo (red) and mIFP driven by *ptc-Gal4* (*ptc-Gal4>mIFP*; *Cherry*:*Smo* /*+*). (G) Same as (F) showing membrane-localized fluorescence only.

Distributions of Cherry fluorescence in Cherry:Smo-expressing discs were essentially mirror reflections of the patterns of GFP fluorescence. Most Cherry fluorescence was close to the apical surface in A compartment cells (Fig. 5E,F). Cherry fluorescence was lower in A cells adjacent to the compartment boundary compared to more anterior cells. In basolateral regions, fluorescence of Cherry:Smo was slightly lower than it was apically, but its apparent presence at the membrane was greater in the P compartment than in the A compartment (Fig. 5F,G). These results are consistent with the idea that degradation of Smo is elevated in cells with lower levels of Hh and that most Smo degradation is apical.

We also examined fluorescence distributions in genotypes that change Hh levels. In *dispatched* (*disp*) mutant discs, Hh transport from P to A compartment cells is compromised (Burke et al., 1999), and we observed that A compartment cells in *disp* mutant discs had lower levels of GFP fluorescence and higher levels of Cherry fluorescence (Fig. 6A,B). In contrast, wing discs that ectopically expressed membrane bound Hh (Hh:CD2) had higher levels of GFP fluorescence and lower levels of Cherry fluorescence (Fig. 6C,D). These results are consistent with the idea that Smo turnover in the A compartment is Hh-dependent.

**Figure 6.**
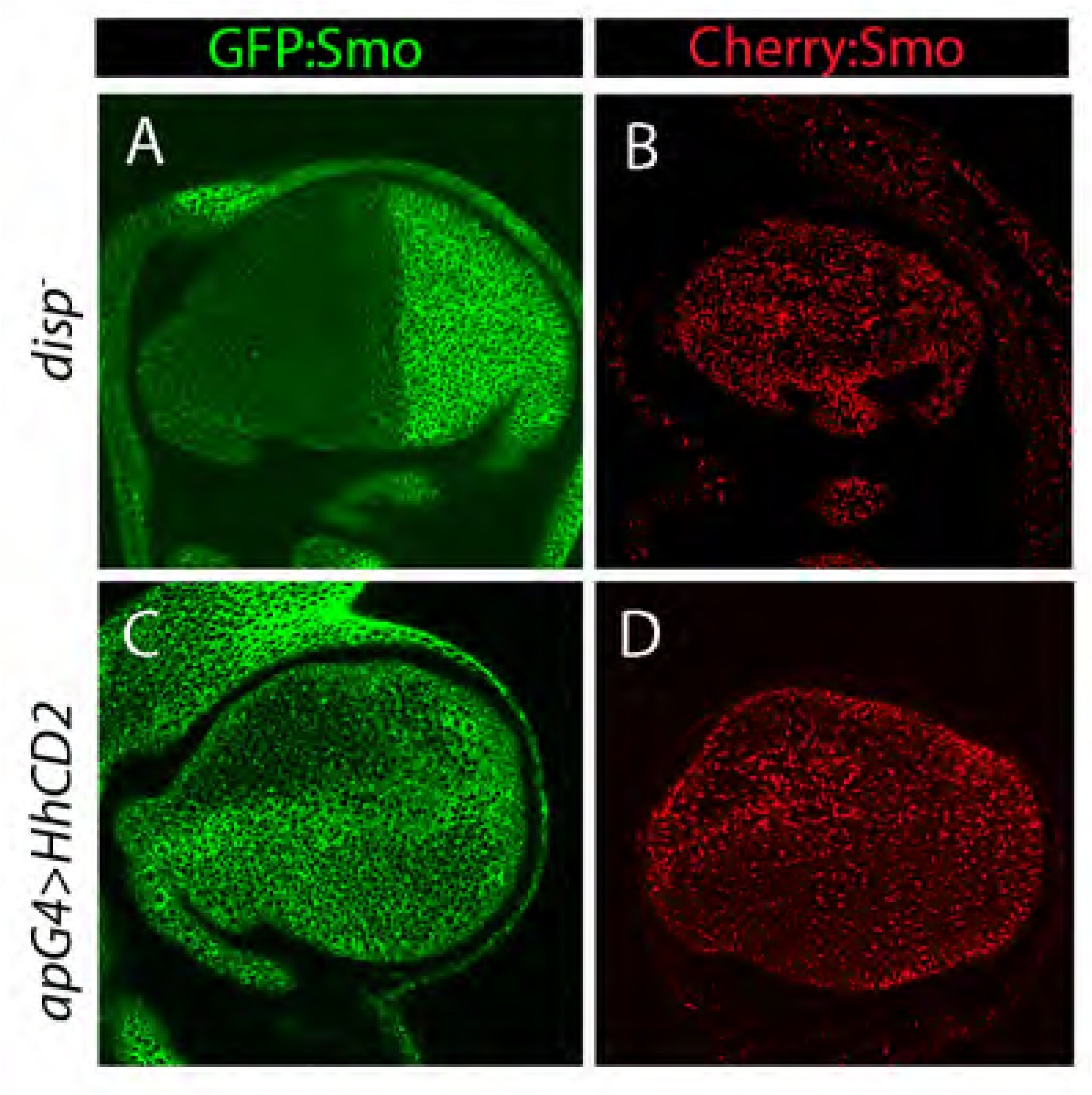
Hh regulation of Smo levels. Frontal views of unfixed wing discs, either expressing BAC-encoded GFP:Smo (green) or Cherry:Smo (red). Images were taken at basolateral optical planes. (A) *disp*^-^ *; GFP:Smo/+* (B) Membrane-tethered Hh:CD2 overexpressed in the dorsal compartment (*ap-Gal4>Hh:CD2; GFP:Smo/+*). (C) *disp*^-^ *; GFP:Smo/+* (D) e Membrane-tethered Hh:CD2 overexpressed in the dorsal compartment (*ap-Gal4>Hh:CD2; Cherry:Smo/+*).

### Measurements of Smo residence time at basolateral membranes

To better understand the kinetics of Smo basolateral to apical movement, we analyzed the maturation of GFP and Cherry fluorescence in the wing disc. To do this, we created a GFP:Cherry tandem fusion protein and performed a pulse-chase experiment in wing discs to measure the time interval between expression of the GFP:Cherry fusion protein and detection of GFP and Cherry fluorescence. We incubated larvae carrying *tub-Gal80^ts^*, *ptc-Gal4*, and *UAS-GFP:Cherry* constructs for one hour at 37°C (the non-permissive temperature for Gal80^ts^) to express the cytoplasmic GFP:Cherry tandem protein, and then acquired images after returning the larvae to permissive temperature (18°C) to stop expression (Fig. 7A-C). GFP fluorescence was first detected approximately 2.5 hours after the heat pulse, but Cherry fluorescence remained almost undetectable (Fig. 7C-E). During the following several hours, both GFP and Cherry fluorescence increased, and the ratio of GFP and Cherry fluorescence plateaued after approximately 8+ hours. The maturation halftimes for GFP and Cherry fluorescence were approximately three and six hours, respectively. The tandem GFP:Cherry fusion protein was distributed uniformly in the cells, suggesting that in contrast to the fluorescent Smo proteins we analyzed, most was not in apical acidic vesicles.

**Figure 7.**
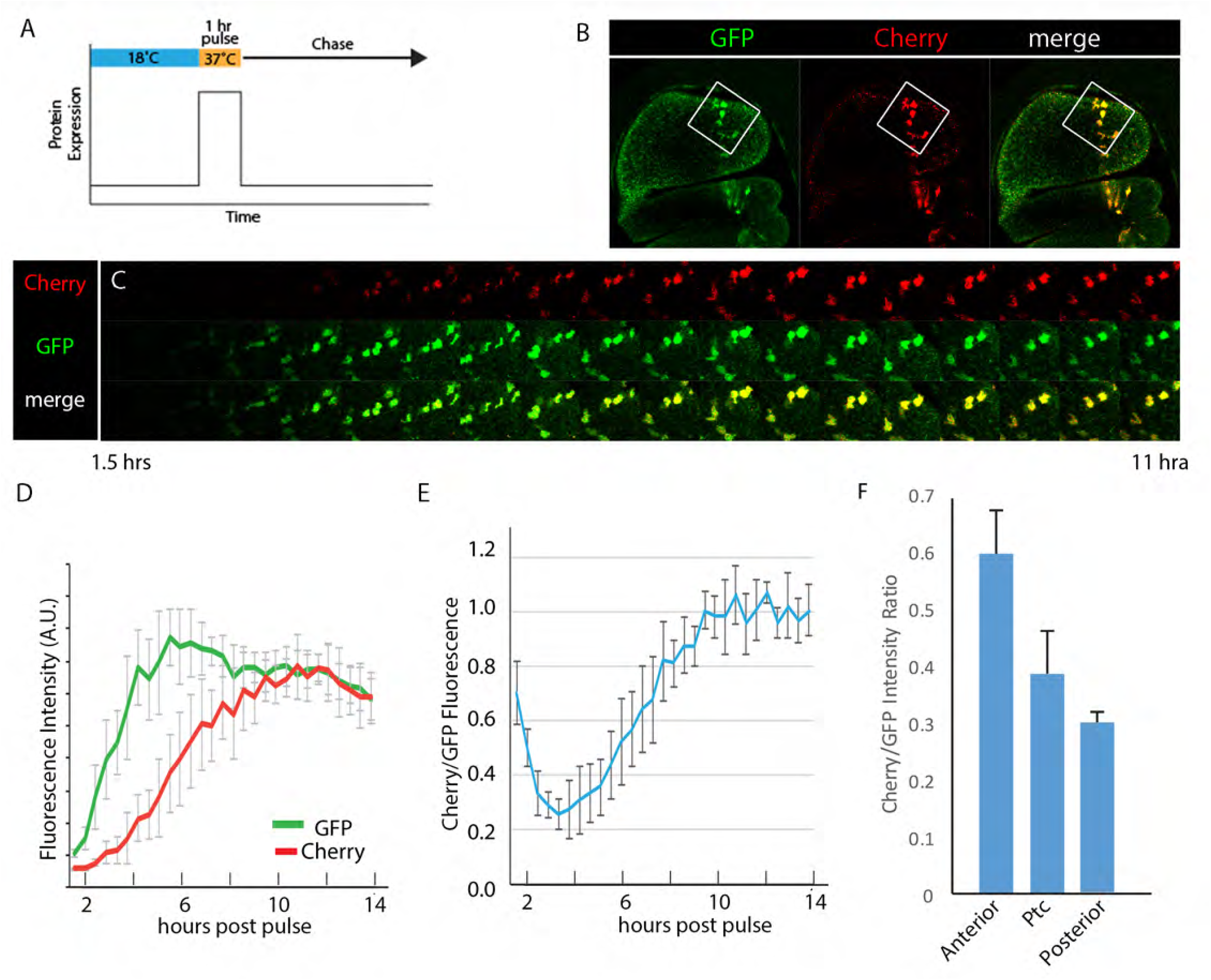
Estimating the residence time of Smo at basolateral membranes. (A) Schematic of the pulse-chase expression of GFP:Cherry tandem timer protein. (B) Frontal sections of unfixed wing discs 12 hours after pulsed expression of GFP:Cherry driven by *ptc-Gal4*. Box indicates cells that are shown at higher magnification in (C). (C) Images of GFP:Cherry fluorescence from cells boxed in (B) at increasing times following expression pulse. (D) Fluorescence of GFP and Cherry following pulsed expression of GFP:Cherry. (E) Ratio of Cherry/GFP fluorescence over time (hours) after pulsed expression of GFP:Cherry. (F) Cherry/GFP fluorescent ratio of BAC encoded GFP:Cherry:Smo in A and P compartments and in Ptc-expression domain.

To relate these measures to estimates of the residence time of Smo at the basolateral membrane, we measured the ratio of GFP to Cherry fluorescence of GFP:Cherry:Smo at the basolateral membrane, and compared this ratio to the maturation kinetics of GFP:Cherry (Fig. 7F). In *ptc*-expressing cells, the residence time was lower than in P compartment cells (<4.5 hours) and lowest in A cells farther from the border (<2 hours). These results suggest that Hh signaling may change the residence time of Smo at basolateral membranes by a factor of two.

### Degradation regulates tissue-level distribution of Smo

Previous studies have reported that the amount of Smo protein is regulated by Hh (Denef et al., 2000). Consistent with this idea, the amount of Cherry:Smo detected by anti-Smo antibody staining in discs expressing Cherry:Smo was higher in cells with high levels of Hh compared to those with low levels (Fig. 8A,B). The distribution of Cherry fluorescence in unfixed discs (Figs. 2,4,5) is consistent with the idea that degradation may control Smo levels. To directly test this idea, we reduced Smo degradation and then monitored GFP:Smo and Cherry:Smo fluorescence. Usp8 is a deubiquitinase that has been shown to remove ubiquitin from Smo and to inhibit Smo internalization and degradation (Xia et al., 2012; Zhou et al., 2018). Upon overexpressing Usp8 in the dorsal wing disc, antibody staining revealed that total Smo increased in the cells that expressed Usp8 (Fig. 8F-K). The increase was specific to basolateral GFP fluorescence and Cherry fluorescence, while apical Cherry fluorescence decreased. These results are in agreement with the idea that degradation is a key mechanism that regulates Smo levels in the wing disc and are consistent with the idea that Smo basolateral localization precedes its transport to degratory apical endosomes.

**Figure 8.**
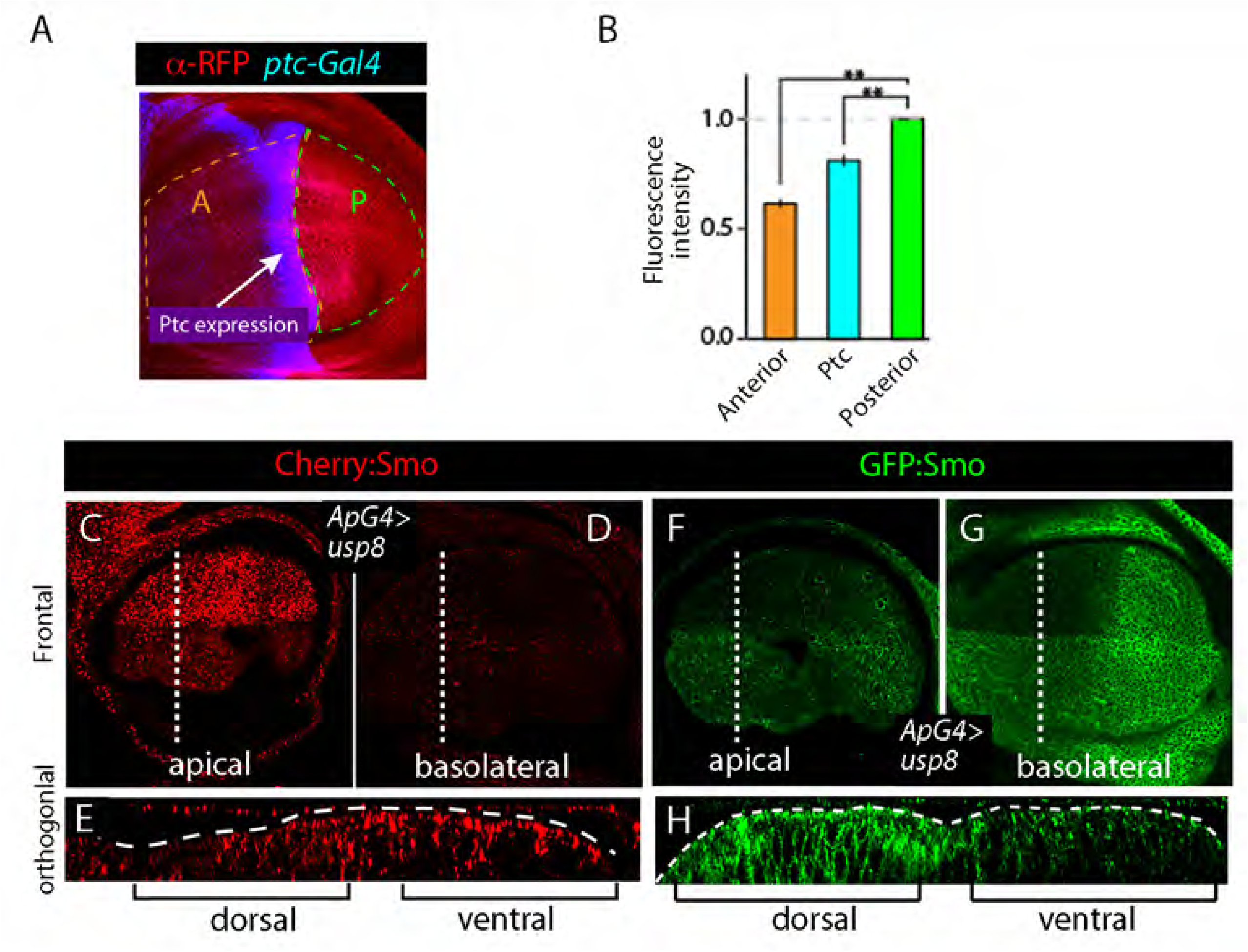
Degradation of Smo controls its tissue distributions. (A) Antibody staining for Cherry (red) and GFP (blue) in wing disc expressing Cherry:Smo and *ptc*-Gal4 UAS-GFP. (B) Quantification of levels of α-RFP fluorescence in A compartment and Ptc expression domain relative to P compartment. (C-E) Wing disc expressing Smo:Cherry (red) in which Usp8 was overexpressed in the dorsal compartment (*ap-Gal4/+; Cherry:Smo /UAS-Usp8*). (C) Apical optical section of frontal view; dotted line indicates location of orthogonal section in (E). (F-H) Wing disc expressing GFP:Cherry (green) in which Usp8 was overexpressed in the dorsal compartment (*ap-Gal4/+; GFP:Smo/UAS-Usp8*). (G) Apical optical section of frontal view; dotted line indicates location of orthogonal section in (H).

### GFP and Cherry fluorescence identifies distinct pools of Smo in embryos and myoblasts

To investigate whether the Smo kinetics and turnover we observed in the wing disc is a general mechanism that also regulates Smo in other tissues, we examined GFP and Cherry fluorescence of GFP:Cherry:Smo in embryos and myoblasts. In stage 11 embryos, Hh is expressed specifically in P compartments and Hh signaling in A compartments is essential for segmentation; and similar to the wing disc, Smo protein levels are elevated in P compartment cells relative to cells in A cells (Alcedo et al., 2000). Antibody staining directed against GFP and Cherry produced striped patterns in GFP:Cherry:Smo-expressing embryos, as expected (Fig. 9A). In live, unfixed embryos, however, GFP fluorescence was similar to the pattern detected by antibody but Cherry fluorescence was more uniform and had no apparent segmental periodicity (Fig. 9B). At the subcellular level, GFP:Cherry:Smo detected by antibodies against GFP and Cherry revealed that most GFP:Cherry:Smo was membrane associated (Fig. 9C). Most GFP fluorescence was also membrane associated in live embryos (Fig. 9D). but Cherry fluorescence in live embryos appeared punctal. These observations are consistent with the observed Smo distributions in the wing disc, suggesting that the fluorescence of GFP and Cherry also identify distinct pools of Smo in the embryo.

**Figure 9.**
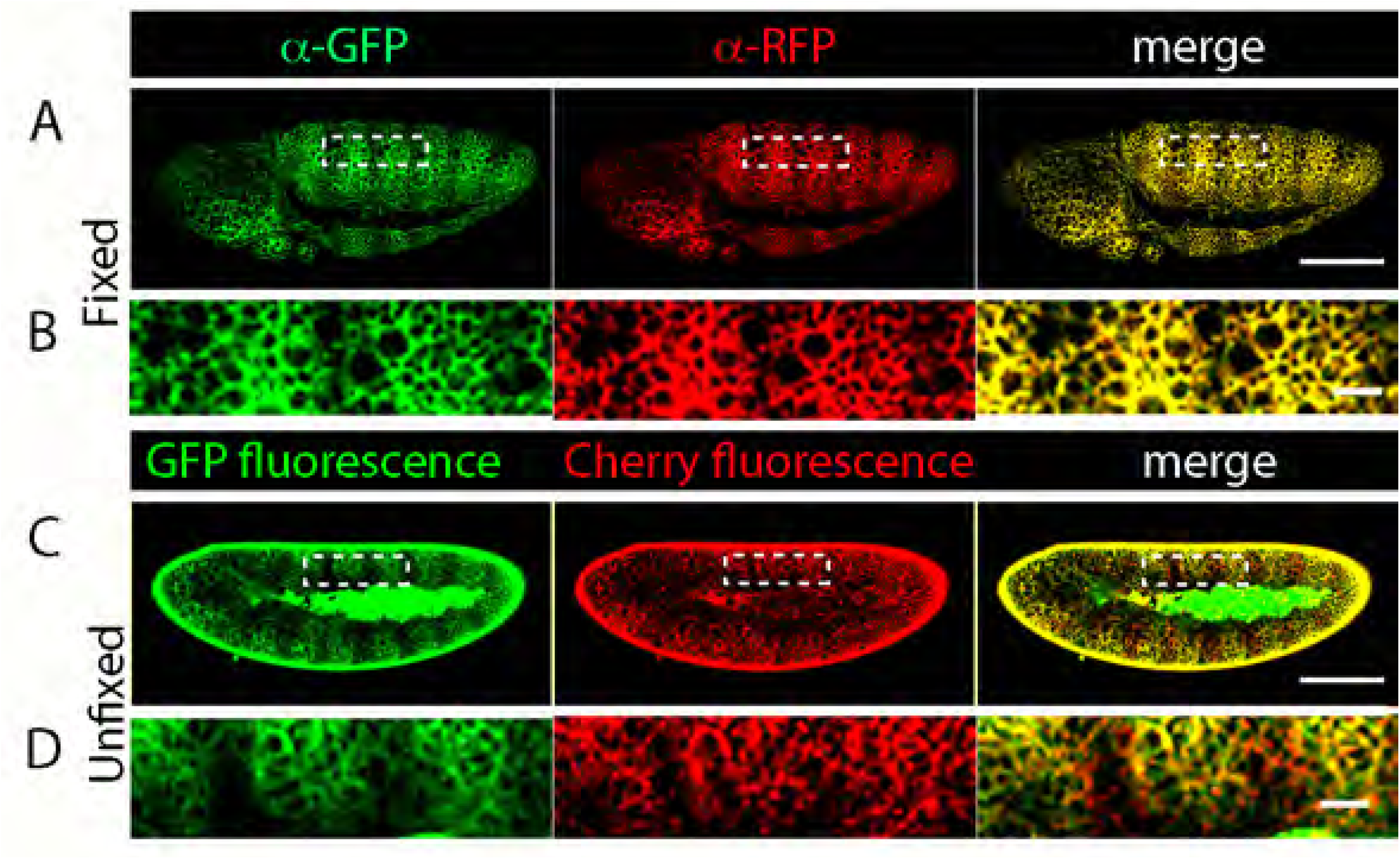
Smo:GFP and Smo:Cherry fluorescence identifies distinct Smo populations in the embryo. (A,B) Stage 12 embryos with BAC-GFP:Cherry:Smo. (A) Fixed embryos stained with α-GFP (green) and α-RFP (red). (B) Higher magnification images of boxed regions in (A). (C,D) GFP (green) and Cherry (red) fluorescence of unfixed stage 12 embryos (C). (D) Higher magnification images of boxed regions in (C). Embryos oriented anterior (left), posterior (right), dorsal (up), ventral (down). Scale

Next, we examined Smo distributions in myoblasts that associate with the larval wing disc but do not have such pronounced apicobasal polarity. Our previous work showed that some of these myoblasts activate Hh signal transduction (Hatori and Kornberg, 2020). First, we tested if the wing disc myoblasts express Hh:GFP encoded by the BAC transgene, but did not detect GFP fluorescence except for sparce puncta in the myoblasts that directly overly Hh-expressing P compartment wing disc cells (Fig. 10). These myoblasts (Fig. 10A,B) have low levels of Hh (Fig. 10C) and higher levels of Ptc and Smo (Fig. 10D,E). When we induced clones of cells that overexpress Smo-RNAi in these cells, we observed a significant reduction of Ptc expression in the clones (Fig. 10F). Together, these results are consistent with the idea that the myoblasts are active for Hh signaling.

**Figure 10.**
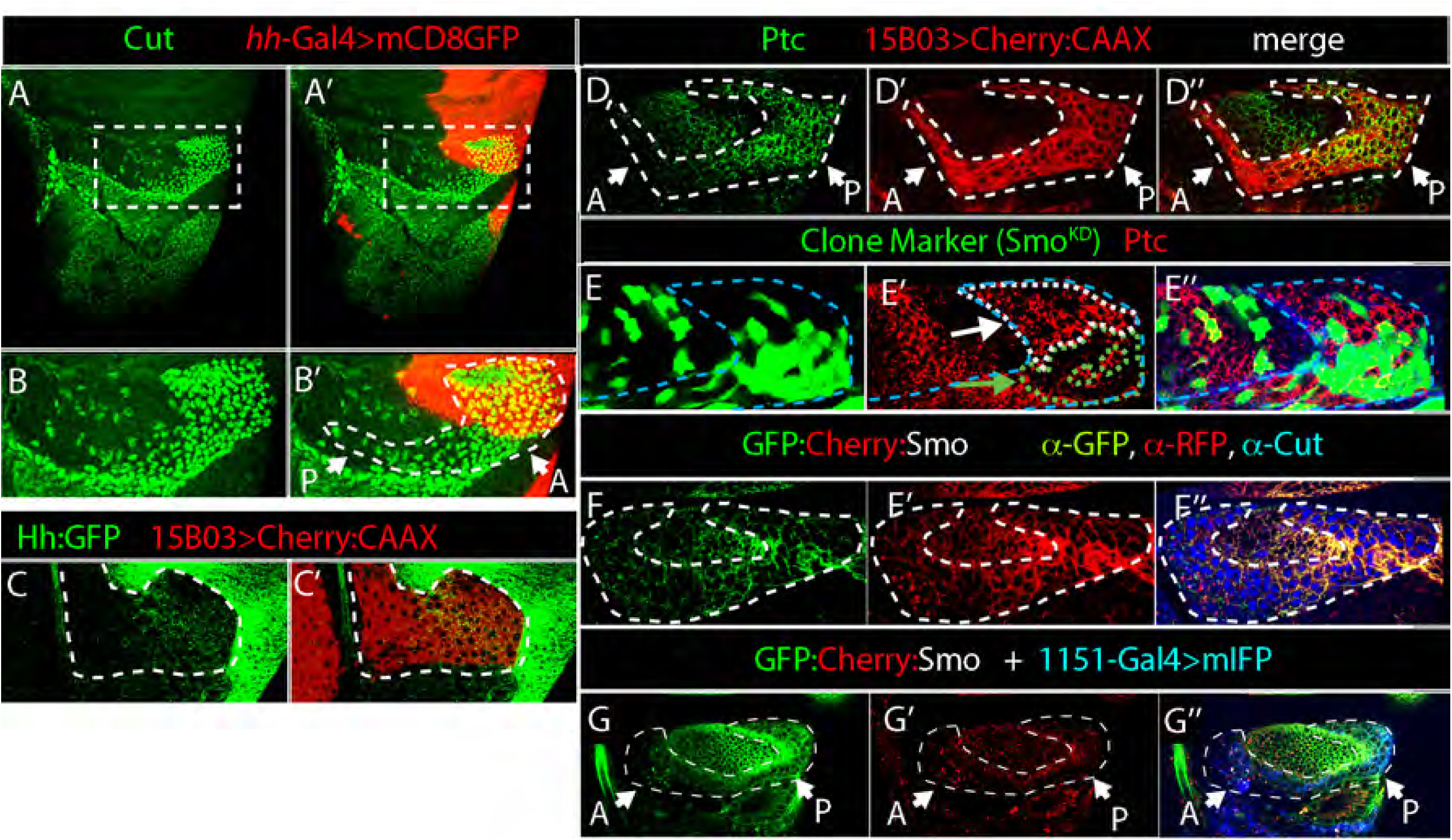
Fluorescence of GFP:Smo and Cherry:Smo in different locations in myoblasts. (A-B’) Wing discs expressing mCD8:Cherry (red) by *hh*-Gal4 stained with α-Cut antibody to mark myoblasts (green). (A) notum region; (B) higher magnification image of boxed region in (A). Dashed white line encircles myoblasts. (C) Distribution of BAC encoded Hh:GFP (green) in the myoblasts (bounded by white dashed line; α-RFP, red). (D) Myoblasts encircled by white dashed line and marked by mCD8:Cherry (15B03Gal4>mCD8:Cherry; α-RFP, red) and α-Ptc antibody staining (green). (E-E”) Myoblasts (outlined with white dashed line and marked by expression of Cut, blue) expressing BAC encoded GFP:Cherry:Smo stained with α-GFP (green) and α-RFP (red) antibodies. (F) Clones induced as in (Rao et al., 2015) expressing SmoRNAi (green) stained for Ptc (α-Ptc, red) bounded by white dashed line. Wild-type myoblasts are bounded by blue dashed line. (G-G”) Myoblasts (outlined with white dashed line and marked by expression of Mef2-Gal4>mIFP, blue) expressing BAC encoded GFP:Cherry:Smo (green) and α-RFP (red).

To monitor Smo regulation in this myoblast population, we also analyzed the fluorescence of BAC encoded GFP:Cherry:Smo. In contrast to the similar patterns of protein detected by anti-GFP and anti-RFP antibodies (Fig. 10E), GFP fluorescence in unfixed discs was higher in more posterior myoblasts compared to more anterior myoblasts, but Cherry fluorescence was uniformly low (Fig. 10G). These results are consistent with the idea that GFP and Cherry fluorescence identify distinct Smo populations.

### Turnover of Cubitus interruptus is slower than that of Smo

We also examined the turnover and kinetics of Ci by using the CRISPR/Cas9 system to add GFP and Cherry tags to endogenously expressed Ci gene (designated as GFP:Ci and Cherry:Ci, respectively). The tags are expected to mark both full length Ci (Ci^FL^) as well as the repressor form (Ci^Rep^) that is generated in the absence of Hh signaling by proteolytic cleavage (Aza-Blanc et al., 1997) (Fig. 11A). *GFP:Ci* and *Cherry:Ci* animals are viable and their wings are wild-type. Trans-heterozygotes with the *Ci^D^* allele (*Ci^D^*/*GFP:Ci* and *Ci^D^*/*Cherry:Ci*) are viable and their wings were indistinguishable from those of *Ci^D^*/+ wings (Fig. 11B-D). We also analyzed the distance between wing veins L3 and L4 in the *Ci^D^*heterozygotes (Fig. 11E-G): the distance between veins L3 and L4 was not distinguishably different in control and mutant flies. These results are consistent with the idea that both GFP:Ci and Cherry:Ci are functional.

**Figure 11.**
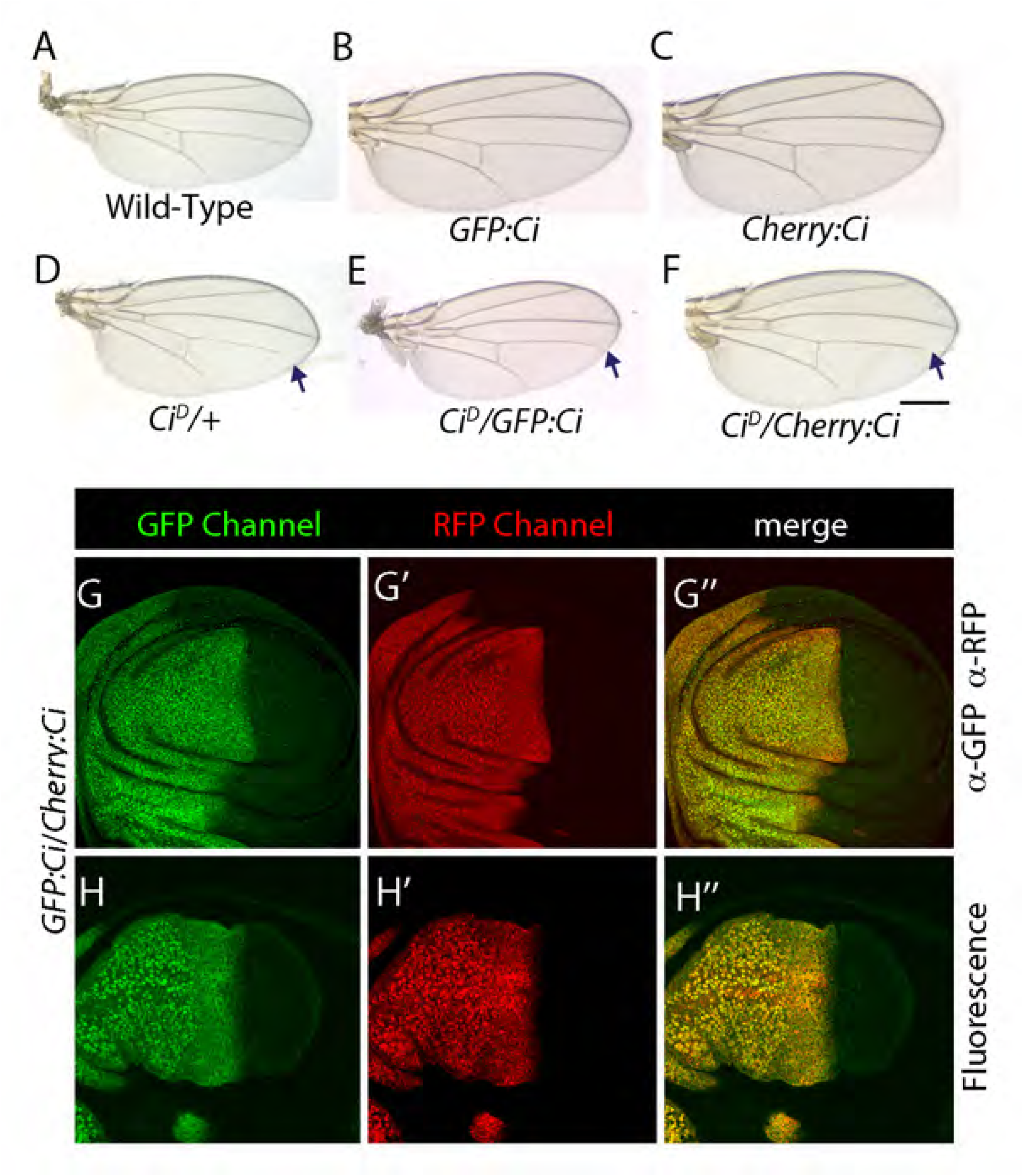
GFP-Cherry timer does not detect Ci turnover. (A-F) Adult wings of indicated genotype. Arrows indicate incomplete vein L4. Scale: 1mm. (G-H”) Optical sections showing frontal views of GFP:*Ci* /Cherry:*Ci* wing discs. GFP (green), RFP (red) Discs were fixed and stained with either α-GFP or α-RFP antibodies (G-G’’) or imaged without fixation (H-H”). (I) Wild-type wing disc stained with α-Ci (AbN). In all images, anterior is to the left, posterior is to the right, dorsal is down and ventral is up.

We compared distributions of GFP:Ci and Cherry:Ci in unfixed wing discs to discs probed with antibodies to GFP or Cherry. Ci detected by the antibodies in fixed discs and Ci fluorescence in unfixed discs were indistinguishable (Fig. 11). These results indicate that kinetics and turnover of Ci could not be distinguished by the different maturation kinetics of GFP and Cherry.

## Discussion

Steady-state distributions of the components of the signal transduction systems which pattern the wing disc provide a framework to understand its development. For instance, levels of Smo, an essential component of Hh signal transduction, are reduced in A cells by Hh-dependent post-translational regulation (Denef et al., 2000). As a consequence, despite uniform *smo* transcription in both A and P compartments, protein levels in the two compartments differ. Another example is the homeodomain-containing transcription factor Engrailed (En), which is exclusively expressed by cells in the P compartment and positively regulates P compartment expression of Hh. Anterior cells express neither En nor Hh, but A cells in a narrow strip adjacent to the compartment border take up Hh from the P compartment and activate signal transduction. These patterns of Smo, En, and Hh protein distribution are necessary for the functionalization of the A cells at the compartment border as a developmental organizer of wing development (Tabata et al., 1995). These patterns appear to be static, but the processes that generate them are not. The work we present here investigated the intracellular distributions of Hh and Smo that contribute to their spatially defined levels in the wing disc.

### Intracellular Hh movements

The discovery that newly synthesized Hh is transported in intracellular vesicles from the apical to the basolateral membrane of P compartment cells (Callejo et al., 2011) was transformative, as it revealed the existence of previously unknown intracellular processes that are essential for Hh signaling. It also made re-evaluation of steady-state distributions imperative, as a particular distribution in a multi-step process can be misinterpreted if the influence of a rate limiting step changes or is not taken into account. For example, the presence of Hh in the apical region of A compartment cells may reflect the rate limiting step in its processing rather than direct movement from the apical membrane of P cells to A cells in the apical lumen. Our finding that approximately 95% of Hh is not apparently released from the P compartment cells that make it (Fig. 1) adds further support for the idea that understanding intracellular movement and regulation of Hh is important (Hatori et al., 2021).

A steep, declining concentration Hh gradient is visible in frontal projections such as shown in Figure 1B, and sagittal sections reveal that this gradient is composed of both basolateral and apical pools (Fig. 1C). Importantly, these images show that the Hh that constitutes this gradient is predominantly if not entirely intracellular. Neither immunohistochemistry nor the fluorescence microscopy we used has the resolution to distinguish whether protein in the apical pool is in the extracellular lumen of the disc or is intracellular and sequestered at or near the apical membrane. Nonetheless, these images of apical Hh show that both Rab7^+^ and Rab7^-^ Hh vesicles are in the same optical plane, consistent with the idea that most (or all) apical Hh is intracellular. Definitive evidence for extracellular Hh in the apical lumen is therefore lacking, in contrast to the strong evidence for Hh in basal cytonemes that move Hh from the P to the A compartment (Chen et al., 2017; González-Méndez et al., 2017),

Although one interpretation might be that apical and basolateral pools are taken up separately from the P compartment (Ayers et al., 2010), we suggest that the Hh we detected in the A compartment is instead consistent with the idea that all the Hh was taken up basally (Chen et al., 2017; González-Méndez et al., 2017) and that basolateral Hh is destined for apical late endosomes. The Hh we detected was punctate or vesicular, and >80% of the apical Hh was in Rab7-containing vesicles (Fig. 1L). These Rab7^+^ vesicles are likely to be late endosomes. This interpretation supports the recycling model of Guerrero (Callejo et al., 2011). Lastly, we note that the images of native Hh:GFP in unfixed preparations show that the large pool of Smo protein in A cells is likely transformed by a relatively small amount of Hh

### Timing intracellular trafficking

The chromophore maturation times for the GFP and Cherry proteins we used to tag Smo differ by approximately three hours (Fig. 7). This temporal disparity made it possible to distinguish between “younger” Smo that was exclusively basolateral and “older” Smo that was exclusively apical (Figs. 4,5). The contrast between these fluorescence distributions was enhanced by the fact that GFP fluorescence, but not Cherry fluorescence, is quenched in an acidic environment, and apical Smo is in acidic, late endosomes (Fig. 3). The images of GFP- and Cherry-tagged Smo show that: 1) Smo moves apically from basolateral locations; 2) that turnover has a significant impact on its overall level and intracellular distribution; and 3) that rates of turnover differ in the A and P compartments.

Smo in the P compartment is both basolateral and apical. Whereas GFP fluorescence of basolateral Smo was bright and membrane-associated, Cherry fluorescence of basolateral Smo was at low levels and punctate (Fig. 5D,F). In contrast, GFP fluorescence of apical Smo was essentially undetectable, and Cherry fluorescence of apical Smo was relatively bright, and punctal. Ptc is a negative regulator of Smo (Chen and Struhl, 1996; Ingham et al., 1991), and Smo is presumably constitutively active in P compartment cells due to the absence of Ptc. The fact that *smo* mutant clones in the P compartment have not been reported to be abnormal (Chen and Struhl, 1996) suggests that Smo activation of the Hh pathway is without effect in P cells, perhaps due to repression of pathway targets by En.

In the A compartment, both α-Smo antibody staining and Cherry:Smo fluorescence was punctal and elevated apically (Fig. 5). GFP fluorescence was uniformly low, suggesting that its half-life is lower than in the P compartment. The higher levels of Cherry:Smo fluorescence in A cells suggests that degradation is rate-limiting, perhaps because the flux of Smo through the processing pathway is greater. Our data do not distinguish whether all Smo cycles through the basolateral membrane and that its residence time in the membrane is regulated by the Hh pathway, or that more Smo is directed to the basolateral membrane by an activated Hh pathway.

## Acknowledgements

This work was funded by NIH R35GM122548 to T.B.K.

## Materials and Methods

### Drosophila lines

The following transgenic lines were used: GFP:Smo BAC (Chen et al., 2017), Cherry:Smo BAC (this study), GFP:Cherry:Smo BAC (this study), Rab5:YFP (Zhang et al., 2007), Rab7:YFP (Zhang et al., 2007), UAS-Rbcn3A RNAi (Bloomington# 34612), UAS-GFP:Cherry (this study), *Ptc-Gal4, ap-Gal4* (Bloomington# 3041), UAS-usp8 (Xia et al., 2012), Cad:GFP (Huang et al., 2011), UAS-smoRNAi (VDRC#9542); Cad:Cherry (Huang et al., 2011), Nrx:GFP fosmid (VDRC# 024184), UAS-CD4:mIFP (Huang et al., 2011; Yu et al., 2015), *P{neoFRT}82B P{law}dispS037707* (Bloomington # 53711) mef2-Gal4 (McQuibban et al., 2010), Cherry:Ci (CRISPR, this study), GFP:Ci (CRISPR, this study), Hh:GFP BAC (Chen et al., 2017), UAS-HhCD2 (Strigini and Cohen, 1997), HS-FLP, UAS-mcd8GFP (Bloomington# 28832); Smo-RNAi (VDRC# 9542), 15B03-lexA (Bloomington# 52486), lexAop-CherryCAAX (K Basler), *smo^3^* (Bloomington# 3277), Tub-Gal80^ts^ (Bloomington# 7018).

### Immunohistochemistry, Fluorescence imaging, and image analysis

L3 larvae were dissected in PBS, and wing discs together with the Tr2 trachea were fixed in PBS+4% formaldehyde. After washing in PBS-TritonX-100 (0.3%), samples were blocked in Roche Blocking Solution. The following antibodies were used: mouse α-GFP (Roche), rabbit α-RFP (Rockland), mouse α-Ptc (DSHB, Apa1), mouse α-En (DSHB, 4D9), DAPI, Phalloidin, mouse α-Dlg1, (DSHB, 4F3), α-Rab7 (Tanaka and Nakamura, 2008), α-Smo (DSHB, 20C6), Alexa633 conjugated-Phalloidin (Invitrogen), DAPI (Invitrogen), AbN (Aza-Blanc et al., 1997). Secondary antibodies from Invitrogen were used and samples were mounted in Vectashield (Vector labs).

Live wing discs were observed using the hanging drop method (Huang and Kornberg, 2016). To neutralize acidic compartments, final concentration of 50mM ammonium chloride was added to the PBS of the hanging drop and samples were images. All images were taken using the FV3000 Olympus Confocal microscope with GaAsP PMT detectors. All images were analyzed and processed on ImageJ and Photoshop. Intensity profiles of confocal pictures were measured using the plot profile function in ImageJ.

### Pulse Chase experiment

Ptc-Gal4, Gal80^ts^/UAS-GFP:Cherry larva were grown at 18˚C to late L3 stage, then incubated at 37˚C for 1 hour, dissected and mounted for imaging. Images were taken every 15 minutes.

### Generation of Cherry:Smo BAC line

Drosophila expressing BAC encoded-Cherry:Smo was engineered using similar methods as described in (Chen et al., 2017).

### Generation of GFP:Cherry:Smo BAC line

Using CRISPR-Cas9, a superfolder-GFP (sfGFP) tag was inserted between the C terminus of Smo and mCherry in line expressing BAC encoded-Smo:Cherry.

#### GFP:Cherry:Smo template donor vector

Three fragments of DNA - left homology arm, GFP tag, and right homology arm - were amplified by PCR. The left homology arm fragment contains an overlapping sequence with PBS-SK vector and mCherry. The Cherry fragment contains an overlapping sequence with the left homology arm and right homology arm. The right homology arm fragment contains an overlapping sequence with mCherry and PBS-SK vector. The three fragments were joined together and cloned into PBS-SK vector using Gibson Assembly (New England Biolabs). The resulting vector is designated as GFP:Cherry:Smo donor vector.

The left arm homology arm was amplified from Wild Type DNA using the following primers. L-arm-fwd: cggtatcgataagcttgatcaccttgccgcacaaatacatatac

L-arm-rev: CCTCGCCCTTGCTCACCATCTCCAGGCCACCGCCCTCTCCGGCAGACACGTCCCGA

The Cherry tag was amplified using the following primers:

Cherry-fwd:TGTCTGCCGGAGAGGGCGGTGGCCTGGAGATGGTGAGCAAGGGCGAGGAGGAT AAC

Cherry-rev:CGCTTGTTCCGGCCGCCCTTCTCTAACTTGTACAGCTCGTCCATGCCGC

The right arm homology arm was amplified from Wild Type DNA using the following primers: R-arm-fwd:GGACGAGCTGTACAAGTTAGAGAAGGGCGGCCGGAACAAGCGGCAGCCGA R-arm-rev:ccgggctgcaggaattcgatGTCATTATTCGGTTATGCTCTCGCTAG

#### pCFD-3 gRNA vector

gRNA sequences were as follows: CGCTCCATTCGGGACGTGTCTGG

The gRNA sequence without the PAM was cloned into pCFD-3 vector obtained from Addgene.

#### GFP:Cherry:Smo fly lines

pCFD-3 vector and donor vector were co-injected into Cas9 expressing flies (nanos-Cas9) by Rainbow Transgenics. The resulting CRISPR-modified flies were screened and verified by sequencing: GFP:Cherry:Smo. The GFP:Cherry:Smo :GFP:Cherry Crispr lines are homozygous viable and transheterozygote of Smo amorphic flies and Smo:GFP:Cherry Crispr flies are also viable.

### Generation of Cherry:Ci and GFP:Ci CRISPR lines

Using similar strategy to generate GFP:Cherry:Smo, the N terminus of the endogenous Ci gene was tagged with either mCherry or sfGFP sequence using CRISPR-Cas9.

The following primers were used:

#### GFP:Ci

L-arm-fwd: cggtatcgataagcttgatGTTTTGCGCTGTTTGTGGACGTTAGAGTG L-arm-rev:

GGATGAGCTGTACAAGTCATGGACTAACTTTAATGAAATGGACGCGTACGCG

GFP fwd: CATTAAAGTTAGTCCATGACTTGTACAGCTCATCCATGCCCAG GFP rev: CGACGTCATTCTTGTTGATGTCCAAGGGCGAGGAGCTGTTCAC

R-arm-fwd: CTCCTCGCCCTTGGACATCAACAAGAATGACGTCGTTTTATAAATTC

R-arm-rev: ccgggctgcaggaattcgatGGCGTTGCCAATAACTTTTGCG

gRNA sequence: AAAATATGTAGGTAACGCGTAGG

#### Cherry:Ci

L-arm-fwd: cggtatcgataagcttgatGTTTTGCGCTGTTTGTGGACGTTAGAGTG L-arm-rev:

GGACGAGCTGTACAAGTCATGGACTAACTTTAATGAAATGGACGCGTACGCG

Cherry fwd: CATTAAAGTTAGTCCATGACTTGTACAGCTCGTCCATGCCGC

Cherry rev: CGACGTCATTCTTGTTGATGGTGAGCAAGGGCGAGGAGGATAAC

R-arm-fwd: CTCGCCCTTGCTCACCATCAACAAGAATGACGTCGTTTTATAAATTC

R-arm-rev: ccgggctgcaggaattcgatGGCGTTGCCAATAACTTTTGCG

gRNA sequence: AAAATATGTAGGTAACGCGTAGG

### Generation of UAS-Cherry:GFP line

sfGFP and mCherry DNA fragments were amplified and were joined with an intervening linker. Thefragment sfGFP-LINKER-mCherry was cloned into pUAST, and pUAST-sfGFP-LINKER-mCherry was injected into Drosophila embryos.

